# “Self-inhibition of HER2 at fever temperatures may prevent dimerization with HER3”

**DOI:** 10.1101/2023.01.22.525115

**Authors:** Puneet K. Singh, Razvan C. Stan

**Author notes:** Corresponding author: Razvan C. Stan - Department of Basic Medical Science, Chonnam National University, Republic of Korea.

## Abstract

**Background:** HER2 receptor is overexpressed in different aggressive and recurrent tumors, and the modulation of its conformational changes has been achieved using immunotherapy or chemotherapy.

**Objective:** To assess the role of fever temperatures on the dynamics of HER2 in the absence of a binding partner.

**Methods:** Molecular dynamics simulations of apo-HER2 were performed at 37□-40□, in order to gauge its intramolecular dynamics.

**Results:** HER2 assumes an inactive and “bent” conformation at 40□, while it maintains its extended, active conformation that can dimerize with other HER partners, in the 37□-39□ range.

**Conclusion:** These novel results indicate the role of fever temperatures on modulating conformational changes in an essential tumor target. Cancer treatment can leverage the use of thermal therapy at particular fever points, in order to complement existing radio or chemotherapy options.

## Introduction

Human epidermal growth factor 2 (HER2) is a ligand-free tyrosine kinase receptor that is a key signaling node in some of the most aggressive tumors, and is an intensely pursued drug target [1]. HER2 differs from other members of the epidermal growth factor receptors (EGFR) family in that it does not bind EGF-like ligands. HER relies instead on hetero-dimerization with other (ligand-bound) EGFR-family receptors for its activation, leading to regulation of cell growth, or, when overexpressed, to involvement in cancer progression. Ligand-bound HER3 is the preferred dimerization partner of HER2 [2]. HER2 ectodomain (ECD) is composed of four domains (I-IV), of which domain II with its “dimerization arm”, a short beta-hairpin comprising residues 243-255, together with a loop of domain IV (residues 571-595) interact and lock with HER3, thus forming an active dimer [3]. Domain IV of the HER family is rather rigid, and, in contrast to the extracellular domains of the other HER receptors, its ECD assumes a fixed conformation resembling a ligand-activated state, permitting it to dimerize in the absence of a ligand [4]. For HER3, there are structural data depicting an auto-inhibited configuration, in which the domain II “dimerization arm” is occluded by intramolecular interactions with domain IV [5]. This intrinsic HER3 auto-inhibition is likely to be overridden in cancers, where HER2 is hyper-activated owing to massive overexpression, although the mechanism through which this is achieved is not elucidated [5]. HER2–HER3 dimer dynamics is preeminent in modulating receptor activity, with recent data linking the strength of EGFR activity with the degree of separation between their domains IV [6].

Here, we asked whether febrile temperatures that are commonly present in infectious diseases and in some types of cancers are relevant for modulating the dynamics of HER2, and in particular, of its domain IV. Using molecular dynamics (MD) simulations, we evidence a mechanism of self-inhibition of HER2 based on the conformational changes of domain IV at 313K (40□), but not at the physiological temperature (310K, 37□) or at mild to acute fever (38-39□). This auto-inhibition may impair the formation of dimers with HER3 and thus attenuate the pro-oncogenic signaling. Although the cancers where HER2 overexpression is present and leads to disease recurrence do not commonly induce a fever response, there is a yet unexplored potential to utilize thermal therapy as an adjuvant to cancer treatment [7, 8].

## Materials & methods

The PDB structure of 6OGE [9] was retrieved from the RCSB database and missing residues were added with the missing residue module of Modeller 10.2 [10]. Simulations were carried out with the Gromacs2020 (http://www.gromacs.org/) using CHARMM force field periodic boundary conditions in the ORACLE server. The structures were solvated in a truncated octahedron box of a simple point charge water model. The solvated system was neutralized with Na+ or Cl− counter ions using the *tleap* program. Particle Mesh Ewald was employed to calculate the long-range electrostatic interactions. The cut-off distance for the long-range van der Waals (VDW) energy term was 12.0□Å. The system was then minimized at maximum force of 1000.0 KJ/mol/nm in 50,000 steps. The solvated and energy minimized systems were than equilibrated for 100 ps under NVT and NPT ensemble processes. The dynamics were integrated using the velocity Verlet integrator, with a time step of 2 fs and bonds constrained using the LINCS algorithm. The simulations were performed for 200 ns at four different temperatures (310K, 311K, 312K and 313K), in triplicates. The results were analyzed using GROMACS toolkits gmx rms, gmx rmsf, and gmx gyrate tools for RMSD, RMSF, and Rg respectively.

## Results & Discussion

A comparison of the structures of monomeric and ligand-bound dimeric ECD of EGFR showed considerable flexibility in the region linking the ectodomain with the transmembrane domain, while mutational studies confirmed the absence of structural rigidity therein [11]. We investigated the role of physiologically relevant temperatures in modulating the dynamics of this region, and of the HER2 as a whole, as shown in Figure 1 for structures obtained at 310K and 313K. For clarity, structures obtained at 311K and 312K are shown in Supplementary Figure 1.

**Figure 1.**
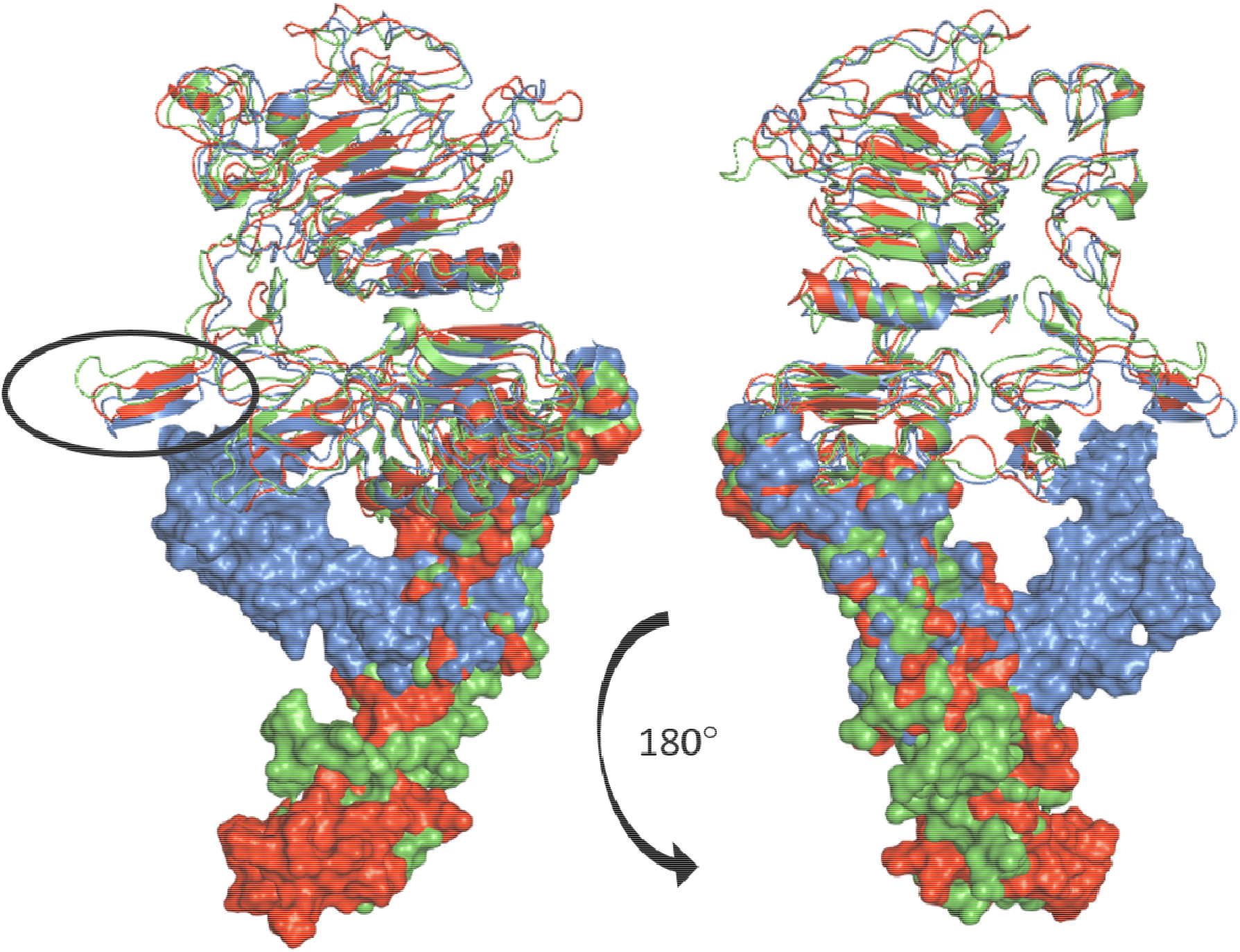
Snapshots taken after 200 ns of HER2 at 310K (red) and 313K (blue). HER2 from PDB 6OGE is shown in green as a reference. Domains IV in all structures are shown as surfaces. The circle indicates the position of the “dimerization arms” of domain II.

In ligand-free states, all EGFR (HER1, HER4 and HER4) are known to adopt a tethered, folded over conformation that is inactive [12], whereas HER2 is an outlier, with an extended arrangement of its domains, as is the case with structures we obtained at 310K-312K. In contrast with the extracellular domains of the other three HER receptors, therefore, HER2 adopts a rigid conformation that allows is to dimerize in the absence of a ligand [4], and that has also been observed in recent structures of HER2-HER3 dimers [13]. This atypical extended state of HER2 is readily accommodated in the functional HER2-HER3 dimers, without the need to undertake additional conformational changes [13]. The structures we observed at 313K are however in a “closed” configuration similar to conformations present in the other ligand-free, inactive HER receptors.

We further investigated the conformational transitions from the native to possible heat-modulated states of HER2, as manifested through global structural parameters, with a focus on the Root Mean Square Deviation (RMSD) and Radii of gyration (Rg) parameters, at different temperatures. We used *R*g to relate the compactness of proteins with changes in temperature during the 200 ns of the simulations (Figure 2.a). *R*g monitors the dimensions of proteins during the simulations and correlates with the rate of folding and the maintenance of protein folds, with lowest *R*g indicating a tighter packing [14]. Calculating RMSD across the simulation time of protein backbones permits the quantification of the degree of conformational changes, as these may occur during MD simulations and can inform on the stability of proteins under different temperatures [15]. An overview of the *R*g and RMSD curves for each temperature point, is shown in Figure 2 and in Supplementary Figure 2.

**Figure 2.**
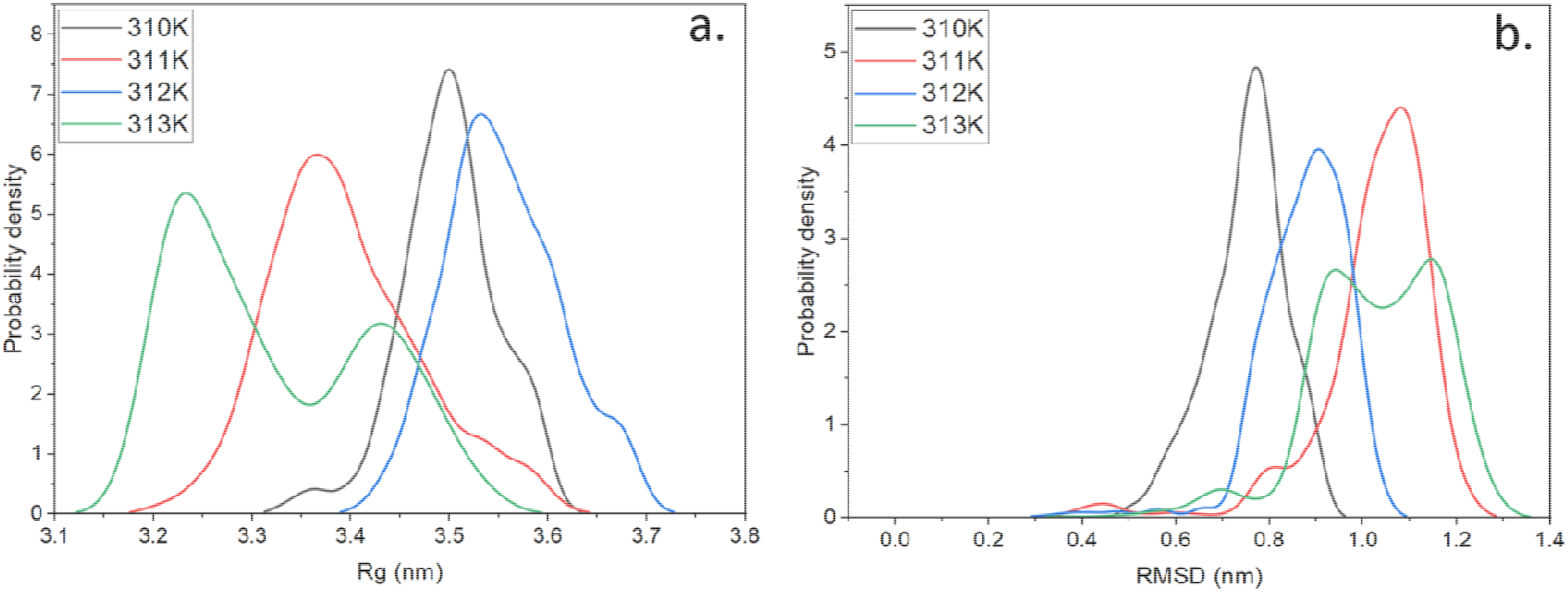
R*g* probability distribution of backbone C_α_ for HER2 at different temperatures (a) and RMSD plots of backbone C_α_ for HER2 (b). Values averaged from three independent trajectories.

The most compact HER2 conformation, as measured using R*g* parameter, is present at 313K. Interestingly, at 313K, a smaller peak is also measured, probably reflecting the movement of domain IV towards domain II that has only been observed at this temperature. Indeed, this movement is not present in any HER3 structures observed after only 100 ns. In turn, the least compact conformation is present at 312K, indicative of subtle conformational changes that can be brought about even by small temperature changes.

RMSD data indicated that HERs is most stable at 310K, with an average centered at 7Å, and becoming progressively less stable with temperature. Intriguingly, two peaks are again observed at 313K for HER2, indicating enhanced mobility that mirrors R*g* data. Of note, the inclusion of domain IV in MD simulations of unbound HER2 has been shown to increase the RMSD of the overall protein [16].

The conformational changes observed in HER2 at 313K versus those present at 310-312K resemble the ligand-induced activation of the αV/β3 integrin complex, which on ligand binding switches from a highly “bent” and inactive conformation, to a more extended, but high-affinity structure with the globular head exposed [17]. That the constitutively extended and active HER2 [18] can be rendered into a folded configuration that is inactive at a fever temperature point may constitute an avenue for cancer therapy. In the clinical literature, there are only a few studies investigating patients with recurring breast cancers, who also presented fever episodes. An investigation of these patients revealed maximal body temperatures that ranged between 38.4 to 39.4□ for up to 20 weeks [19], along with metastasis to liver. HER2 itself is also present in hepatocellular carcinoma patients with liver metastasis that present a fever response stemming from the pyrogens released by the necrotic tissue [20]. For breast cancer patients in stage III (with infiltrating tumors without metastases), hyperthermia in a temperature range of 40 to 43□ for about one hour is routinely used [21]. Targeting HER2 has been successfully achieved with thermosensitive liposomes containing chemotherapeutic drugs, and guided by membrane embedded HER2 targeting antibody (Herceptin), but only after the tumor temperature has been raised to 40 to 41□ [22]. Fever-range temperatures may also affect HER2 clustering on membranes that are known to be involved in its signaling [23].

## Conclusions

We determined that only at 40□, HER2 receptor adopts a conformation that may render it unable to form productive dimers with other HER partners, and may lead to a diminished role in tumorigenesis. While a core body temperature of 313K is not achieved clinically during cancer progression, means to leverage our observations through the judicious use of thermal therapy may be relevant for the treatment of pertinent cancers.

## Conflict of interests

The authors report there are no competing interests to declare.

Human and Animal Rights Not applicable.

## Acknowledgments

Funding was provided by National Research Foundation of Korea, grant 2021R1I1A2059587 (RCS). This work was supported by Oracle Cloud credits and related resources provided by the Oracle for Research program.

## Notes

### Competing Interest Statement

The authors have declared no competing interest.

