## Supplementary Figure 1 for "“Self-inhibition of HER2 at fever temperatures may prevent dimerization with HER3”"

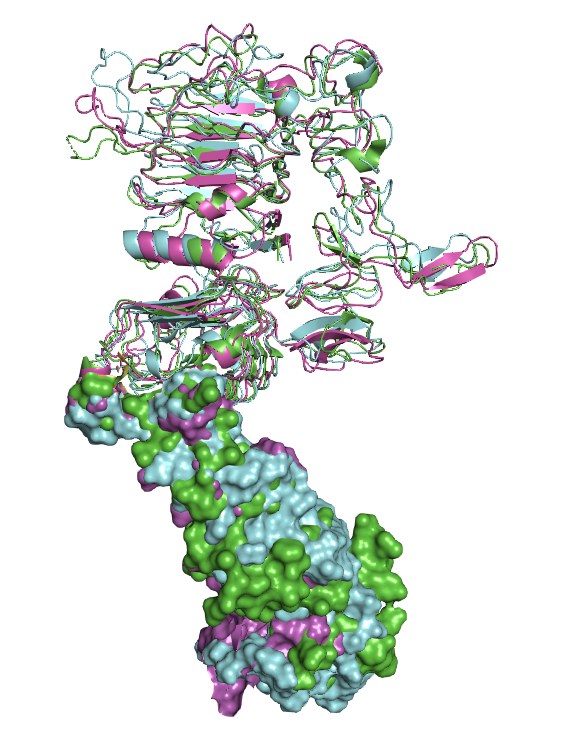

Supporting Figure 1. Snapshots taken after 200 ns of HER2 at 311K (magenta) and 312K (cyan). HER2 from PDB 6OGE is shown in green as a reference. Domains IV in all structures are shown as surfaces.
